# Can Deep Convolutional Neural Networks Learn Same-Different Relations?

**DOI:** 10.1101/2021.04.06.438551

**Authors:** Guillermo Puebla, Jeffrey S. Bowers

## Abstract

Same-different visual reasoning is a basic skill central to abstract combinatorial thought. This fact has lead neural networks researchers to test same-different classification on deep convolutional neural networks (DCNNs), which has resulted in a controversy regarding whether this skill is within the capacity of these models. However, most tests of same-different classification rely on testing on images that come from the same pixel-level distribution as the testing images, yielding the results inconclusive. In this study we tested relational same-different reasoning DCNNs. In a series of simulations we show that DCNNs are capable of visual same-different classification, but only when the test images are similar to the training images at the pixel-level. In contrast, even when there are only subtle differences between the testing and training images, the performance of DCNNs could drop to chance levels. This is true even when DCNNs’ training regime included a wide distribution of images or when they were trained in a multi-task setup in which training included an additional relational task with test images from the same pixel-level distribution.

## Introduction

Relational reasoning is core to human intelligence (Penn, Holyoak, & Povinelli, 2008), and has proven to be a challenge for an earlier generation of connectionist models (e.g., O’Reilly & Busby, 2002; Rogers & McClelland, 2004; St. John, 1992) and more recent deep networks as well (Ricci, Cadène, & Serre, 2021; Vankov & Bowers, 2020; Puebla, Martin, & Doumas, 2021). Perhaps the simplest form of relation reasoning is the same-different task that simply requires the reasoner to determine whether two inputs are the same or different by some criterion. In the domain of vision, the simplest version of this is to classify images as visually identical or not. This skill is essential to abstract combinatorial thought and seems to be much more developed humans and chimpanzees than in other species (Gentner, Shao, Simms, & Hespos, 2021).

Recently there has been mixed evidence regarding whether standard deep convolutional neural networks (DCNNs) can support same-different matching of images. Fleuret et al. (2011) developed a Synthetic Visual Reasoning Test (SVRT) that included a set of 23 classification problems involving images of randomly generated shapes (for example images see Fig. 1 “Original” column). They reported that standard machine learning techniques at the time did poorly on same-different tasks. Similarly, Stabinger, Rodríguez-Sánchez, and Piater (2016) showed that state-of-the-art DCNNs (at the time) LeNet and GoogLeNet performed poorly on the same SVRT same-different tasks, and more recently, Kim et al. (2018) showed that vanilla DCNNs were poor at SVRT same-different tasks, and using a different dataset, showed that Santoro et al. (2017) relational network (RN) also failed to support same-different judgments.

**Figure 1:**
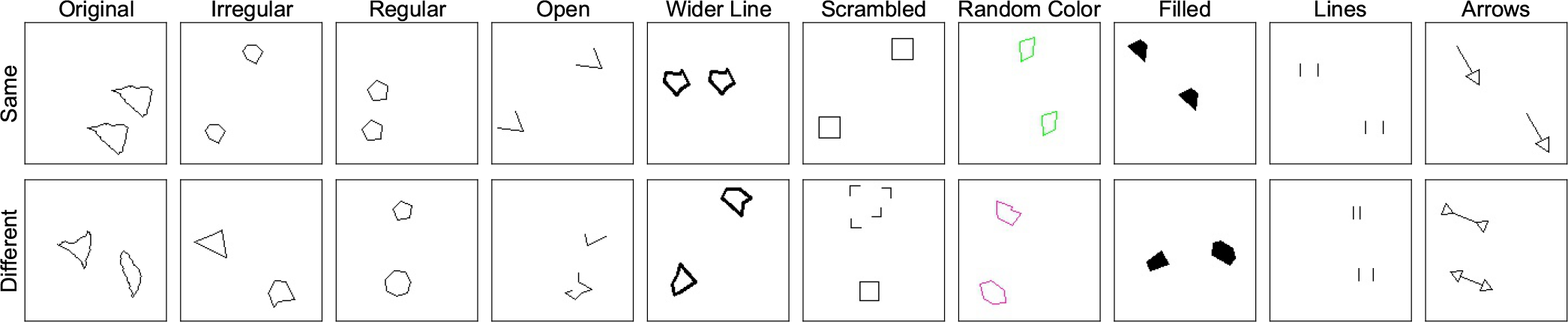
Positive and negative examples from SVRT problem #1 and our new nine versions of this problem.

Interestingly, Kim, Ricci, and Serre (2018) did find that a “Siamese network” that encoded the two images in two separate channels in order to simulate the effects of attentional selection and perceptual grouping learned to classify images as “same” or “different” easily, leading the authors to conclude that object individuation is a key step in solving the same-different task. At the same time, they also argue that a full solution to the same-different problem requires a network to encode dynamic representations of relations rather than statically storing visual-relation templates in synaptic weights. That is, in their view, symbolic processes need to be implemented to fully solve the same-different task.

On the other hand, there are recent reports that the current state-of-the-art DCNNs can solve the same-different task. If this is indeed the case, it would be a striking example of standard networks solving a fundamental relational reasoning task without implementing any symbolic machinery. Funke et al. (2021) noted that Kim et al. (2018) only tested relatively small CNNs (up to 6 layers), and when they replicated the same-different experiments on the SVRT using a ResNet-50 (He, Zhang, Ren, & Sun, 2016) model (a network of 50 layers) the models were able to perform the task successfully. Funke et al. (2021) noted that the success does not necessarily imply DCNNs can solve all visual reasoning tasks, but they do highlight that standard feedforward processing DC-NNs can solve the same-different task and that Kim et al.’s claim regarding the need for extra mechanisms for this task is unwarranted.

Similarly, Messina, Amato, Carrara, Gennaro, and Falchi (2021) have shown that a range of recent DCNNs, specifically ResNets, DenseNets, and CorNet-S, can solve the same-different SVRT tasks, whereas they confirm that this is difficult for older AlexNet and VGG networks. The authors conclude: “We think that the development of the abstract and relational abilities of neural networks is an important leap towards achieving some interesting new tasks…”.

However, there is a fundamental problem with using success on the same-different SVRT task as evidence that CNNs can support relational reasoning. A key feature of relational reasoning is that it is reasoning based on relations rather than any low-level visual details of the inputs. In the domain of visual same-different judgements, reasoning should extend to novel images. The SVRT task does test models on novel pairs of images, but the test images are generated in the same way (i.e., the train and test datasets come from the same pixel-level distribution), and accordingly, it does not test the hypothesis that models have acquired the capacity to support relational reasoning on the same-different task.

## Simulations

In the simulations below we test abstract same-different reasoning in DCNNs based on the ResNet-50 architecture. The basic tenet of our simulations is that a model that has learned the abstract *same* and *different* relations should be able to recognize examples of these relations beyond its training set.

Our training and test data are based on problem #1 of the SVRT (see Fig. 1 column “Original”). In problem #1 images of two randomly generated shapes are classified as “same” if they are the same up to translation and “different” otherwise. Furthermore, we created nine new datasets that followed the same abstract rule as problem #1. However, each new dataset was generated through a distinct stochastic generative process (i.e., a different pixel-level distribution). In the *irregular* dataset each shape was a (randomly generated) irregular polygon. In the *regular* dataset each shape was a regular polygon. In the *open* dataset each shape was an irregular polygon where the first and last vertices were not connected. The *wider line* dataset was the same as the irregular dataset except that the line width was set to two pixels instead of one. The *scrambled* dataset was the same as the regular dataset except that in the “different” examples one the of the objects (scrambled) was generated by dividing the other object into sections and displacing them randomly around the center. The *random color* dataset was the same as the irregular dataset except that for each image the line color was chosen randomly. The *filled* dataset was the same as the irregular dataset except that the shapes were filled with black. In the *lines* dataset each object corresponded to two unconnected vertical lines; in the “same” examples the distances between the lines of each object were exactly the same, whereas in the “different” examples these distances were different. Finally, in the *arrows* dataset the objects were arrows consisting of one or two triangular head(s) and a line; the head(s) and the line were connected; in the “same” examples the arrows were the same and in the “different” examples the orientation of each head was inverted. Note that among these nine different stimulus sets there are differences in the level of low-level similarity with the original SVRT data. In particular, the irregular, regular, and, to a lesser extent, the open datasets are more similar to the original data than the rest of the datasets.

### Simulation 1

In Simulation 1 we created four models based on ResNet-50. All models consisted on a ResNet-50 convolutional front end followed by a hidden layer with 1024 units with ReLU activation (see Fig. 2). In Simulation 1 there was one output layer that consisted of a single sigmoid unit which predicted whether the input image belonged to the category “same”. We pre-trained the models’ convolutional front end in either ImageNet (Deng et al., 2009) or TU-Berlin (Eitz, Hays, & Alexa, 2012), a dataset of human-generated sketches. Furthermore, we varied how we treated the output of the convolutional front end before passing it to the hidden layer. We either applied a global average pooling (GAP) operation to it, as Funke et al. (2021) did, or flatten it, as Messina et al. (2021) did^1^.

**Figure 2:**
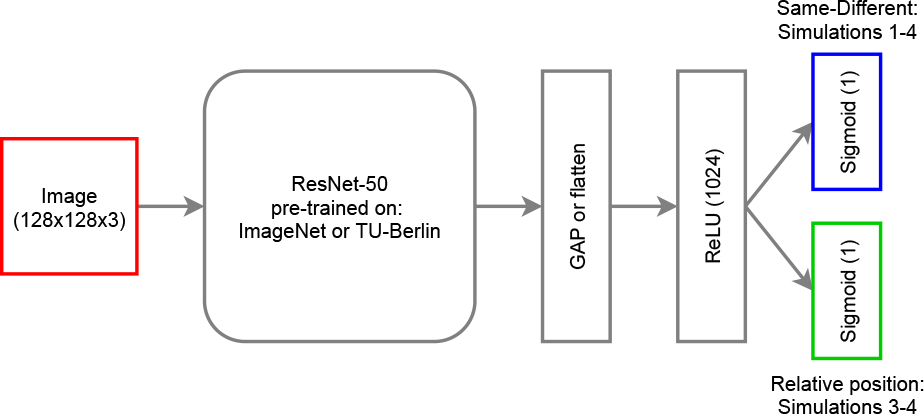
Model set-up.

We trained each model 10 times with the Adam optimizer (Kingma & Ba, 2014). Training proceeded in two stages. In the first stage, the pre-trained ResNet-50 network was frozen while the rest of the network was trained with a learning rate of 0.0003. In the second stage, the complete model was trained with a learning rate of 0.0001. The training data consisted of the original data of the SVRT problem #1. In the first stage the model was trained on 28000 images for 5 epochs with a batches of 64 samples. In the second stage the model was trained on the same images for 10 epochs and with the same batch size.

In Simulation 1 we performed the most basic and stringent test of abstract relational reasoning. The models were trained on problem #1, using the original dataset, and then presented with 5600 images from each of the 10 stimulus sets. That is, our testing conditions consisted on new images from the original training set (replicating Funke et al., 2021), and novel images from the other nine test datasets that were not seen during training. As noted above, a model that has learn the abstract *same* and *different* relations should generalize learning on the same-different task independently from the pixel-level similarity to original SVRT data.

Because we are testing binary classification models on datasets that come from different distributions than the training data, it is possible that each test dataset has a different optimal classification threshold. To account for this, we used the area under an ROC curve (AUC), which is an aggregate measure of performance across all possible classification thresholds. AUC values range from 0 to 1, where 0.5 corresponds to chance-level prediction. As customary, an *AUC* ≤ 0.6 was considered fail, an 0.6 < *AUC* ≤ 0.7 was considered poor, an 0.7 < *AUC* ≤ 0.8 was considered fair, an 0.8 < *AUC* ≤ 0.9 was considered good, and *AUC* > 0.9 was considered excellent.

#### Results and Discussion

As can be seen in Fig. 3, all models achieved excellent performance in the original test dataset. Furthermore, the models pre-trained on ImageNet performed better than the models pre-trained on TU-Berlin and the models with GAP performed better than the models that flattened the last convolutional layer’s output. Overall, the ImageNet & GAP model was the best performing model in the original test dataset as well as across the nine new test datasets. Accordingly, the following analysis (as well as Simulations 2-4) will concentrate on it. The ImageNet & GAP model showed good or excellent performance in the irregular, regular and open conditions. As can be appreciated in Fig. 1, these conditions were the most featurally similar to the training data. On the other hand, the ImageNet & GAP model performed poorly or worse on the random color, lines, and arrows conditions with fair performance on the wider line and scrambled conditions. In general, these results show that the degree of generalization on the same-different task depends heavily on the pixel-level similarity between the training data and the test data. This pattern of results is inconsistent with the models learning the abstract *same* and *different* relations.

**Figure 3:**
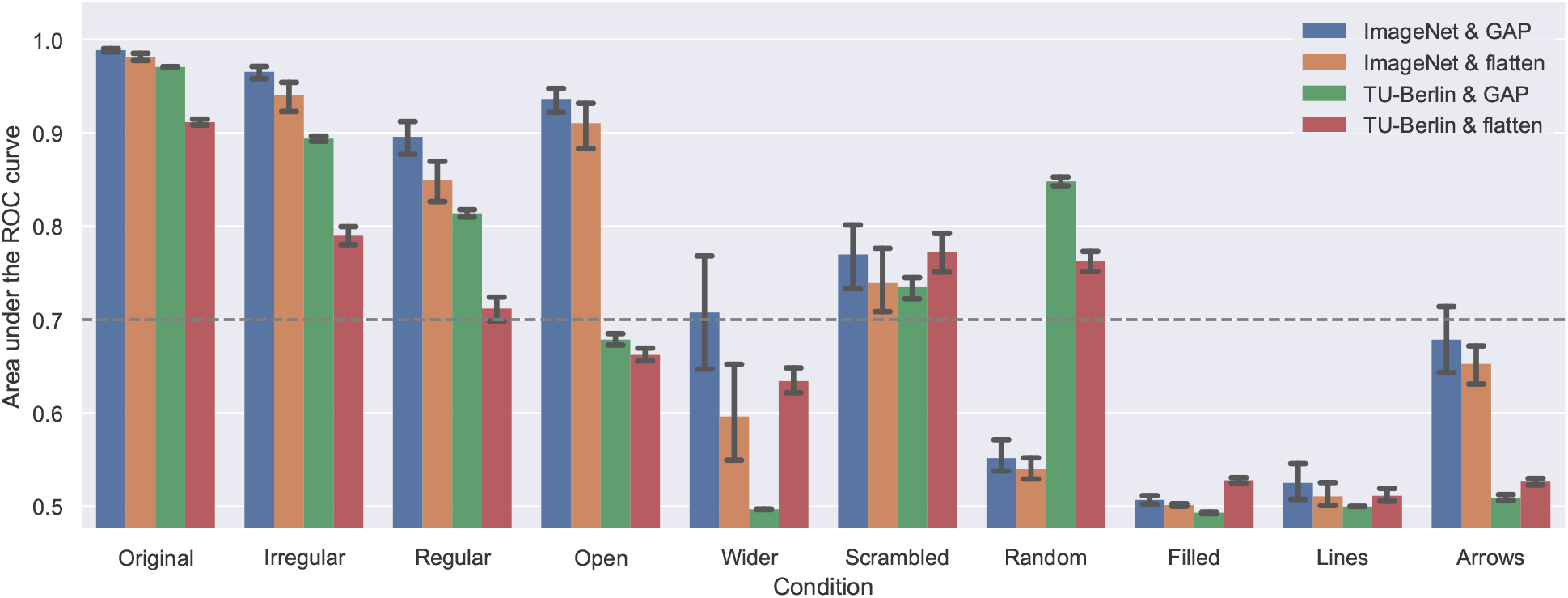
Average AUC by condition and model on 10 runs of Simulation 1. Error bars are 95% confidence intervals.

### Simulation 2

A potential criticism to Simulation 1 is that the training data (line drawings of random shapes) wasn’t rich enough for the models to form a more complex representation of the “same” and “different” relations. Note, however, that Messina et al. (2021) do interpret their results following the same training conditions as supporting abstract same-different reasoning in DCNNs, while Funke et al. (2021) argues that they results suggest that purely feed-forward mechanisms are sufficient to perform abstract same-different visual reasoning. Nevertheless, we agree that it is important to test what happens when the models have access to a richer training set. Therefore, in Simulations 2-4 we tested whether augmenting the training regime of the ResNet-50-based models would improve generalization on the same-different task to unseen stimuli. In Simulation 2, we did this by training the models on nine stimulus conditions consisting of images from the original SVRT data and all the new datasets except one. For each condition we trained 10 models with the same settings as in Simulation 1 except that the models were trained for 13 epochs instead of 10. We tested the models on the one stimulus set they were not trained on. For example, the models in the irregular stimulus condition were trained on the original data and all the new datasets except the irregular condition, on which they were tested.

#### Results and Discussion

As can be seen in Fig. 4, the ImageNet & GAP model showed good or excellent performance in the irregular, regular, open, wider line, scrambled and filled conditions. Presumably, the improvement on the wider line and filled conditions is due to their pair-wise similarity. In contrast, the random color, lines, and arrows conditions were less affected by the additional training, since they were the most featurally unique among the different datasets. These results show clearly that augmenting the training regime of the model with data from several conditions increases the model’s ability to generalize to an untrained condition. However, this benefit seems more related to the pixel-level similarity of the augmented data with the testing data than to the shared relational structure of the problem among conditions. If the model was learning the same/different relation, then the benefits of our data augmentation manipulation would have spread evenly across conditions.

**Figure 4:**
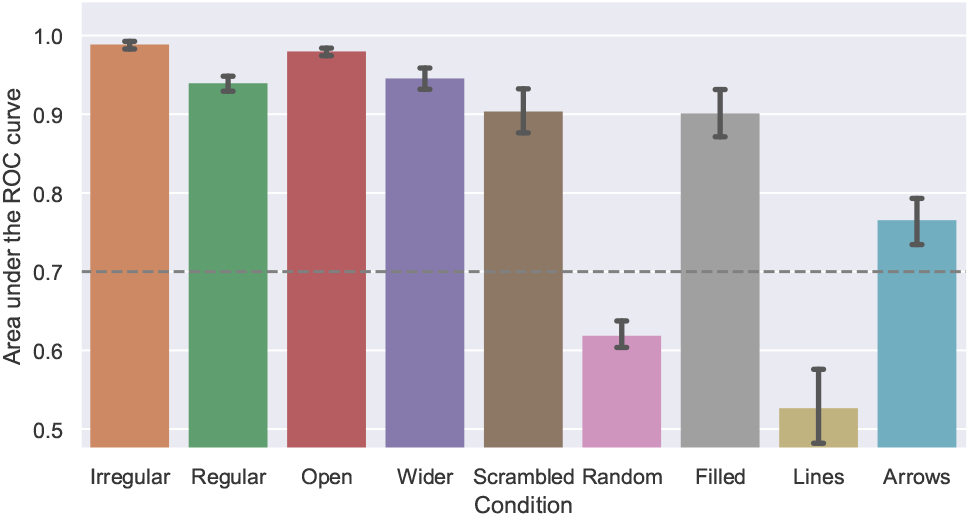
Average AUC by condition on 10 runs of Simulation 2. Error bars are 95% confidence intervals.

### Simulation 3

In Simulation 2 we augmented the models’ experience of the different conditions by training on the same-different task directly. A potential problem with this strategy is that it does not give the models any experience with the specific condition they are tested on. In contrast, in Simulation 3 we augmented the models’ experience on all the datasets through multitask learning. In deep learning research multi-task learning has long been used as technique to improve generalization (Ruder, 2017). In this simulation the models were trained on two tasks. The first was the same-different task as in the previous simulations. The second was a relative position task. This consisted in classifying whether the lower object in the image was to the right of the upper object (category 1) or to the left (category 0). To do this we added a second output layer with a single sigmoid unit (see Fig. 2). Note that the processing path of this architecture only diverges at the output layer. This kind of hard parameter sharing is known to reduce overfitting (Baxter, 1997), so if our previous results are just a matter of overfitting to the training data^2^, adding a second task should aid to generalize learning of the same-different task.

We trained 10 models with images from the “same” and “different” categories from all conditions. However, we only allowed the models to learn to classify images from the original SVRT data as “same” or “different”, whereas the models learned to classify all presented images into their corresponding relative position category. To accomplish this, during training we used the following composed loss function:

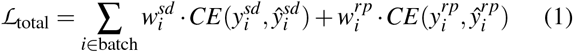

where *CE*(*y, ŷ*) is the cross-entropy loss between the label *y* and the prediction *ŷ*, and *w*^*sd*^ and *w*^*rp*^ are the weights for the same-different loss and the relative position loss, respectively. During training, *w*^*rp*^ was set to 1 for all images. In contrast, when the model received images from the original SVRT data we set *w*^*sd*^ to 1, otherwise it was set to 0. During testing, we presented the models with images of each problem version and recorded the models’ same-different and relative position AUC. All other training and testing parameters were the same as in Simulation 1 except that we trained the models for 13 epochs rather than 10.

#### Results and Discussion

As can be seen in Fig. 5, the ImageNet & GAP model achieved perfect performance in the relative position task. In the same-different task, on the other hand, the results were more varied. The majority of the conditions showed a good or excellent performance except for the filled, lines and arrows conditions. Notably, the lines and arrows conditions did not seem to be affected by training on the secondary relative position task (compare Fig. 5 with Fig. 3). Overall, these results show that training in a “source” task (relative position) can improve generalization on a “target” task (same-different) for out of distribution samples as long as those samples come from a distribution of images trained on the source task. Is important to note that this technique involves to effectively show all the pixel-level distributions of unfamiliar images to the model. Furthermore, our results show that multitask training does not affect all conditions equally. Notably, this technique was less effective for the more featurally unique conditions, suggesting that the similarity between the train data and the test data plays an important role in multitask-learning too.

**Figure 5:**
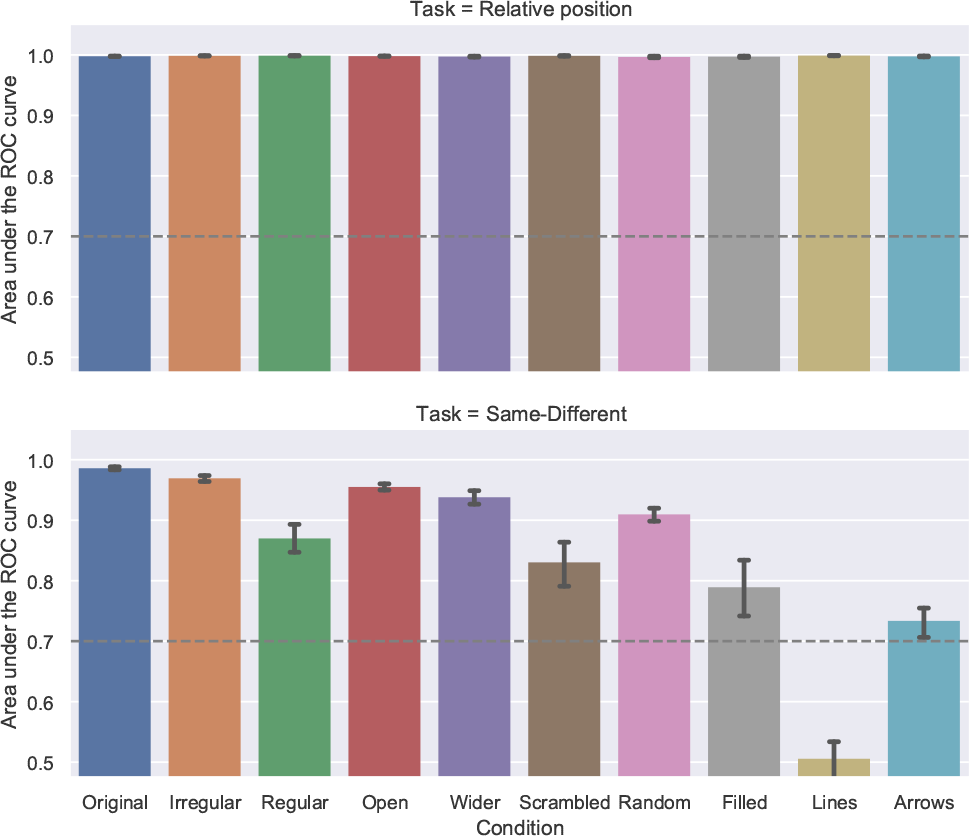
Average AUC by condition and task on 10 runs of Simulation 3. Error bars are 95% confidence intervals.

### Simulation 4

In Simulation 4 we combined the approaches taken in Simulations 2 and 3 in order to provide the models with the maximum amount of information to generalize the same-different task to the unseen conditions. As in Simulation 3, we trained the models in both the same-different and the relative position tasks. Furthermore, as in Simulation 2, for the same-different task we trained on all the stimulus conditions except one. For each of these 9 conditions we trained 10 models and tested them on the stimulus set that was not trained on. We trained the models with loss (1), this time setting *w*^*sd*^ to 1 for all datasets except the one tested on. All other training parameters were the same as in Simulation 3.

#### Results and Discussion

As can be seen in Fig. 6, for the relative position task the ImageNet & GAP model achieved perfect performance in all conditions, just as in Simulation 3. The models’ performance in the same-different task was better than in Simulation 3, with the models achieving excellent performance in all conditions except lines and arrows. The cumulative effect of the extra training and secondary task manipulations suggest that our interpretation of the effect of training on a secondary task is indeed akin to a form of data augmentation. Note that these forms of data augmentation improve performance by increasing the similarity of the training data to the test data. This conclusion is supported by the results obtained in the lines and arrows conditions, the most featurally unique conditions in our simulations. The fact that after applying both, extra training on the same-different task on all other conditions and training on all conditions in the relative position task barely affects the models’ performance on these conditions shows that the performance benefits of these manipulations come from the pixel-level similarity of the (heavily augmented) data with the test data. This is not consistent with the models grasping the abstract relational concepts “same” and “different”.

**Figure 6:**
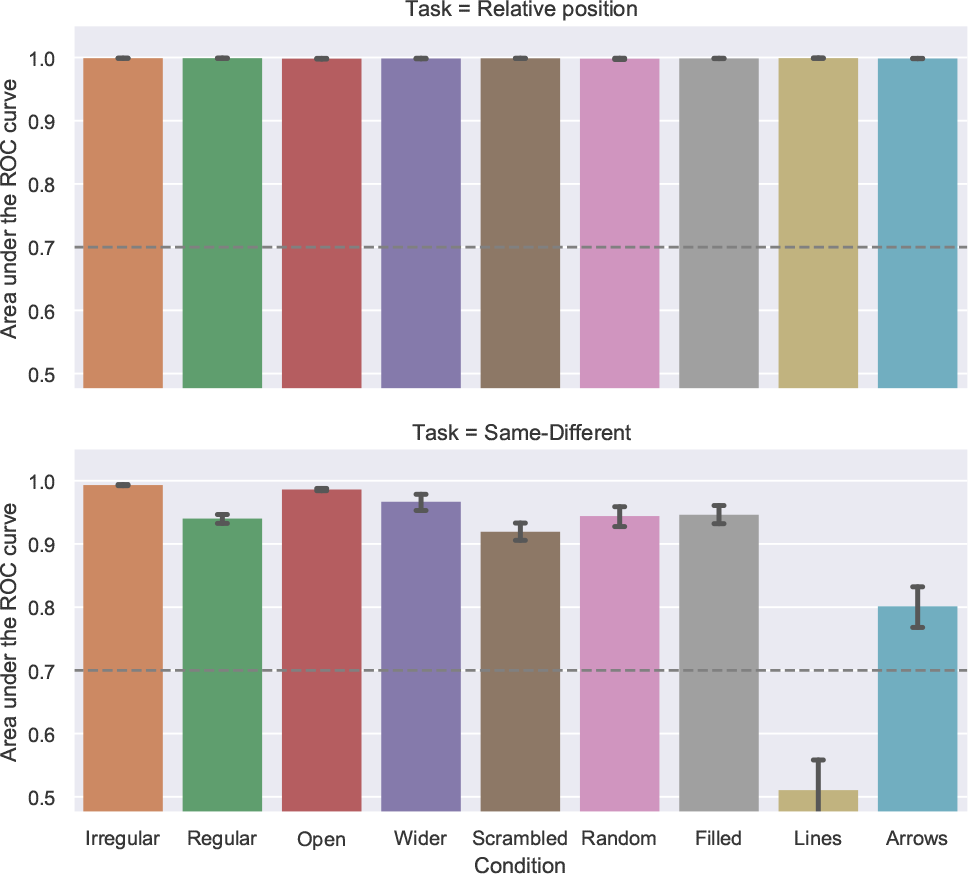
Average AUC by condition and task on 10 runs of Simulation 4. Error bars are 95% confidence intervals.

## General Discussion

In four simulations we tested whether DCCNs were able to learn the abstract *same* and *different* relations that would support relational reasoning in the same-different task. Across simulations we found that, instead of forming an abstract representation of this task, DCCNs were unable to generalize to new test images that shared the same underlying relations as the training data but were not similar at the pixel level. This was the case even when we augmented DCCNs’ experience with new stimulus sets that instantiated the same-different task with several kinds of objects (Simulations 2 and 4), and when we used multi-task learning to give them experience with the very same stimulus conditions that they were tested on (Simulations 3 and 4).

Across simulations we found that, instead of forming an abstract representation of this task, DCCNs generalized primarily based on the pixel-level similarity of the training data with the test data. In contrast, DCCNs did not generalize well to new test images that shared the same underlying relations as the training data but were not similar at the pixel level. This was the case even when we augmented DCCNs’ experience with new stimulus sets that instantiated the same-different task with several kinds of objects (Simulations 2 and 4), and when we used multi-task learning to give them experience with the very same conditions that they were tested on (Simulations 3 and 4).

These results shed new light into the discussion of whether is necessary to invoke extra, symbolic mechanism to solve the same-different task. If by “solving” the same-different task one means generalizing from one set of images to another set of images that share the same pixel-level distribution (as Funke et al., 2021, assume and is implemented in the SVRT test) it is perfectly reasonable to say that DCNNs are able to solve this task. This, by itself, is an interesting problem from a machine learning point of view, because early machine learning models could not solve this kind of task. However, if by “solving” the same-different task one means to learn a representation of the *same* and *different* relations that support generalization beyond pixel-level similarity (as in humans and chimpanzees), our results suggest that DCCNs are just not up to the task.

In future work, we plan to extend the present analyses to relation networks (Santoro et al., 2017). Relation networks are an interesting test case because they are feed-forward neural networks that are specifically designed to perform relational reasoning. However, the way they have been benchmarked so far does not allow to test directly whether their performance is based on low-level similarity between the training and test data or on more abstract representations. The current results suggest that dynamic representations of relations and objects might be necessary to achieve true visual relational reasoning.

## Footnotes

1 We also made models that had the pre-trained convolutional front end frozen and only the classifier was trainable. Those models achieved similar results to the ones presented on Simulation 1. However, they were not well-suited for the data augmentation and multi-task learning techniques used on Simulations 2-4, so we don’t consider them further.

2 Note, however, that the test data of the SVRT problem #1 consists of a different set of images from the training data, so overfitting would have resulted in a low AUC in the original condition, which is the opposite to what we found in Simulation 1.

